# Staying in control: characterising the mechanisms underlying cognitive control in high and low arousal states

**DOI:** 10.1101/2023.04.17.536548

**Authors:** Clara Alameda, Chiara Avancini, Daniel Sanabria, Tristan A. Bekinschtein, Andrés Canales-Johnson, Luis F. Ciria

## Abstract

Throughout the day, humans show natural fluctuations in arousal that impact cognitive function. To study the behavioural dynamics of cognitive control during high and low arousal states, healthy participants performed an auditory conflict task during high-intensity physical exercise (*N* = 39) or drowsiness (*N* = 33). In line with the pre-registered hypotheses, conflict and conflict adaptation effects were preserved during both altered arousal states. Overall task performance was markedly poorer during low arousal, but not for high arousal. Modelling behavioural dynamics with drift-diffusion analysis revealed evidence accumulation and non-decision time decelerated, and decisional boundaries became wider during low arousal, whereas high arousal was unexpectedly associated with a decrease in the interference of task-irrelevant information processing. These findings show how arousal differentially modulates cognitive control at both sides of normal alertness, and further validates drowsiness and physical exercise as key experimental models to disentangle the interaction between physiological fluctuations on cognitive dynamics.

## 1. Introduction

Rushing to get to work on time or falling asleep while watching a TV series are familiar scenarios that illustrate how people’s level of arousal is endogenously altered. These fluctuations in arousal occur throughout the day as a function of circadian rhythms, sleep pressure or physical activity. Ranging from deep sleep to intense physical exertion, arousal fluctuations modulate cognition and information processing (Goupil & Bekinschtein, 2012; Lambourne & Tomporowski, 2010). Here, we investigate how cognitive control is exerted across the arousal spectrum by instructing healthy participants to perform an auditory Simon task (Simon & Rudell, 1967) while transitioning to sleep (i.e., drowsiness) or physical extenuation (i.e., high-intensity physical exercise).

During reduced arousal states such as drowsiness, individuals systematically show poorer overall task performance, i.e., longer reaction times (RTs) and lower accuracy (Bareham et al., 2015), (Xu et al., 2023). Similarly, increasing arousal level with acute exercise close to (or exceeding) the anaerobic threshold seems to raise RTs (González Fernández et al., 2017). For this reason, the interaction between arousal and cognition has been frequently approached from the experimental framework initially proposed by Yerkes-Dodson (1908). According to their well-known inverted U-shaped theory, an intermediate level of arousal will be associated with optimal cognitive performance, while levels above or below will lead to a decrease in performance, as recently found in drowsy and high-intensity exercising participants (Ciria et al., 2021) for probabilistic reversal learning, a higher order cognitive task.

Of particular interest are the studies manipulating arousal levels to investigate cognitive control. Cognitive control is the ability that allows us to adaptively adjust cognitive processes so that task-relevant information is maintained and non-relevant or conflicting information is inhibited (Botvinick & Braver, 2015). This ability is commonly assessed by means of conflict tasks such as the Simon task (Simon & Rudell, 1967), the Stroop task (Stroop, 1935) or the Flanker task (Eriksen, 1995), in which two main cognitive control-related effects can be measured: 1) cognitive conflict (i.e., longer RT in incongruent vs. congruent trials); and 2) conflict adaptation (i.e., reduced conflict effect when the previous trial is incongruent vs. congruent). Regarding the effect of high arousal on cognitive control, the existing literature suggests that stimulus-response conflict effect is still present when performing a conflict task during moderate exercise (Davranche et al., 2015; Davranche & McMorris, 2009). However, the effect of acute exercise on overall RTs in conflict tasks is rather ambiguous: while some studies report shorter RTs (e.g., Alves et al., 2012), others show no effect or even the reverse pattern (e.g., Labelle et al., 2013; Weng et al., 2015). At the opposite side of the arousal spectrum, cognitive conflict effects seem to be also overall preserved in sleep-deprived and drowsy participants (Bratzke et al., 2012; Canales-Johnson et al., 2020; Gevers et al., 2015). Indeed, Canales-Johnson and collaborators (2020) presented an auditory Simon task to participants while they transitioned towards sleep. Stimulus-response conflict and adaptation effects were found to remain unaffected during low arousal, although, as expected, RTs increased.

In this study, we designed a complementary experiment to that of Canales-Johnson et al. (2020) to characterise cognitive performance at both sides of the arousal spectrum by implementing the same auditory Simon task to healthy participants in a state of high arousal (i.e., high-intensity physical exercise). Following the approach by Ciria and colleagues (2021), databases from both experiments were combined aiming to reveal not only whether drowsy and highly aroused participants perform reliably different, but why they do so (i.e., which specific cognitive elements might be modulated by such states). To this end, we applied a drift-diffusion model (DDM) for conflict tasks (Ulrich et al., 2015) to disentangle different elements involved in decision-making and cognitive control processes.

According to DDMs, binary decisions can be captured by a process of accumulating evidence for one answer over the other until a decision threshold is reached, triggering the motor response (Ratcliff et al., 2016). Based on this assumption, DDMs allow to parametrise specific decision-making features, such as rate of evidence accumulation, separation between decision thresholds or non-decision time (i.e., sensory encoding and motor response). Additionally, the specific DDM for conflict tasks (Ulrich et al., 2015) implemented in this study distinguishes between task-irrelevant information (i.e., automatic) processing and task-relevant (i.e., controlled) processing, enabling to also unravel the amount of interference produced by the processing of irrelevant or conflicting information. Previous studies involving conventional DDMs show that drowsy and sleep-deprived states lead to a lower drift rate, wider boundary separation and longer non-decisional time (Jagannathan et al., 2022; Ratcliff & Van Dongen, 2011), while moderate intensity exercise seem to fail to modulate decision-making parameters (Lefferts et al., 2016, 2019).

Based on the premises that (1) drowsiness and high intensity physical exercise both seem to lead to slower RTs and lower accuracy (Goupil & Bekinschtein, 2012; Lambourne & Tomporowski, 2010), and (2) changes in arousal have no apparent impact on conflict and adaptation effects (Canales-Johnson et al., 2020; Davranche et al., 2015); we expected the magnitude of cognitive conflict effects to be preserved in high and low arousal states, although both states would result in higher overall RTs and lower accuracy relative to their respective baseline conditions. Consistent with recent accounts for perceptual decision making (Jagannathan et al., 2022), we additionally expected DDM analyses to reveal a slower rate of evidence accumulation and a greater separation of the decisional boundaries in the low arousal condition. Further, comparisons between high arousal and its baseline were of a broader exploratory nature. All hypotheses here tested, together with the analysis plan, were pre-registered after data collection (https://aspredicted.org/x8qd6.pdf), except for the DDM analyses.

## 2. Materials and methods

### Participants

A total sample of 72 healthy participants (aged from 18 to 50 years old) was included in the present study. All participants reported normal or corrected-to-normal vision and binaural hearing and had no recent history of cardiovascular, musculoskeletal, neurological or psychiatric illness. Before the experimental sessions, all participants were informed of the experimental protocol and signed an informed consent form. They were given monetary compensation for their participation. The studies were approved by the local committee of ethics of the University of Cambridge (CPREC.2015; Experiment 1) and the University of Granada (978/CEIH/2019; Experiment 2).

Experiment 1 consisted of 33 participants (18 females; age range, 18–30) of an initial cohort of 41, pruned after rejecting those who did not attain enough drowsiness for single participant comparisons (i.e., participants with less than 10 trials in a single condition, e.g., less than 10 congruent trials preceded by an incongruent trial in drowsy state^1^). All participants were recruited from the city of Cambridge. Only non-pathological easy sleepers, with a sleepiness score 7-14 as assessed by the Epworth Sleepiness Scale (Johns, 1991), were selected to increase the probability that participants fell asleep under laboratory conditions. They were instructed to get a regular night’s rest on the night before testing, and to avoid stimulants (e.g., coffee/tea) on the day of the experiment.

Experiment 2 consisted of 39 participants (2 females; age range, 18-50) recruited from Granada through announcements on mailing lists and social media. Although a total of 40 participants were initially recruited, data from one participant was discarded, as the heart rate (HR) achieved in the high intensity condition was considered too low, i.e., average HR below 70% of their maximum HR (over three standard deviations below the group average of 85% of HR max [SD = 5.06]). Only physically fit cyclists with a training regime of at least 4 times or more per week and at least 5 years cycling experience were selected, since the execution of a 30-minute task while cycling at 80% of VO2 max requires participants with a high level of training to ensure the safe and proficient completion of the strenuous experimental protocol within the laboratory setting. In addition, testing expert cyclists ensures proper EEG signal-to-noise ratio, since they are used to maintain a fixed posture on the bike for long periods of time and their head movements during pedalling are less abrupt, compared to people who are not used to cycling (Ciria et al., 2018). Participants were asked to get a regular rest the night before each experimental session, and not to engage in strenuous physical activity in the 48 hours prior to the sessions.

### Experimental task

Participants performed a modified auditory version of the Simon task (Simon & Rudell, 1967) in English —in Experiment 1— or Spanish —in Experiment 2—. In this task, recorded spoken words (“left”/”izquierda” or “right”/”derecha”) were presented to either the right or the left ear through earphones. Trials were considered congruent when the word meaning corresponded to the side from which they were heard, and incongruent when they did not. Participants were asked to press with their thumbs the button located on the side which was congruent with the meaning of the word depicted by the voice, while ignoring the physical location from which the word was presented. For example, they should press the left button when they heard the word “left”, even if it was presented through the right earphone (see Figure 1A). In Experiment 1, two response boxes were placed at hand level on a reclined chair. In Experiment 2, response buttons were placed on both sides of the bike’s handlebar. All four types of trials were presented equally often in a random order. The time interval between the participant’s response and the presentation of the following stimulus ranged between 2 and 2.5 seconds in Experiment 1, and between 1.5 and 2 seconds in Experiment 2. The shorter inter-trial interval in Experiment 2 was aimed at increasing the number of completed trials in a shorter total task time. Participants were allowed to respond within 1.99 seconds after the word.

**Figure 1:**
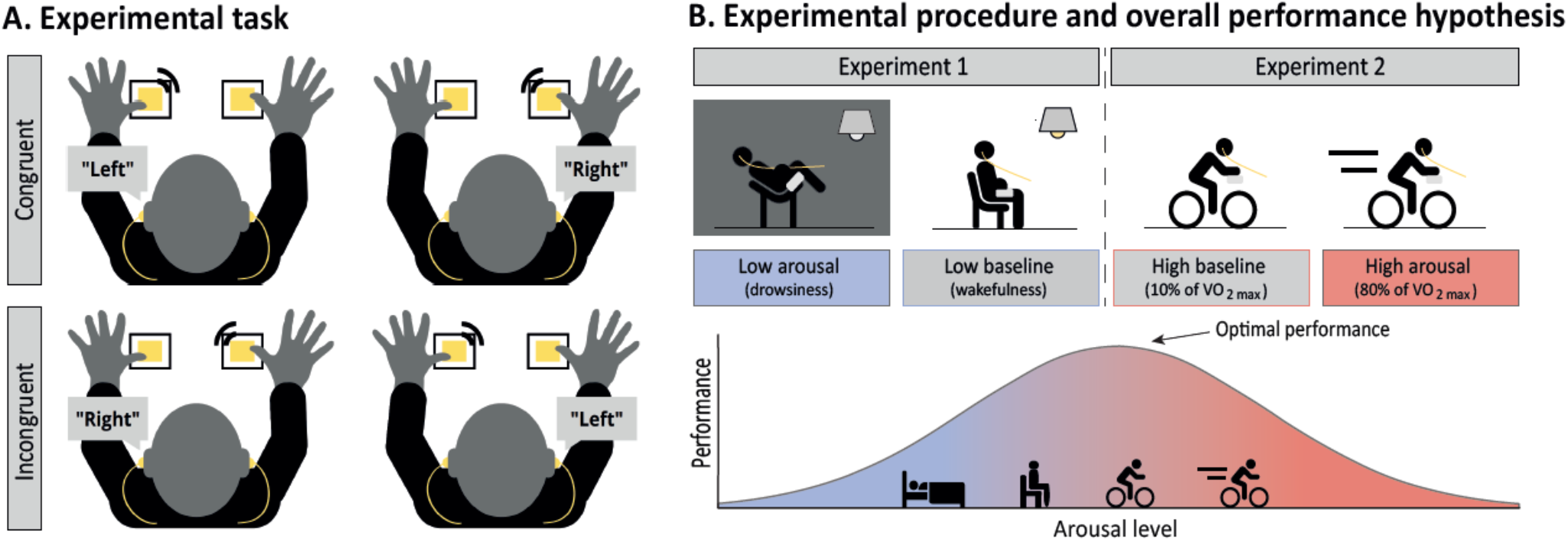
Experimental design. A) Schematic representation of the auditory Simon task implemented. Trials were considered congruent when the meaning of the word matched the ear through which it was presented, and incongruent when it did not. Participants were asked to press the button corresponding to the meaning of the word, ignoring the location in which it was heard. B) Schematic representation of experimental procedure and main hypothesis on overall performance. In Experiment 1, the experimental session was divided into two parts: wakefulness and drowsiness. Experiment 2 consisted of two experimental sessions in which participants performed the task while pedalling at high or low intensity. Optimal performance (i.e., overall RT and accuracy irrespective of trial type) was expected at a moderate arousal state (e.g. exercising at low intensity), while lower (drowsiness) and higher (exercising at high intensity) arousal states were expected to result in poorer task performance (longer RT and lower accuracy).

### Procedure

Throughout two different experiments, we presented the auditory Simon task to participants who were either in states of (1) drowsiness (hereafter referred to as “low arousal”), compared to attentive wakefulness (“low baseline”), or (2) high-intensity physical exercise (“high arousal”), compared to low-intensity (“high baseline”).

### Experiment 1

Experiment 1 consisted of a single experimental session. Participants were fitted with an electrolyte 129-channel cap (Electrical Geodesics, USA) while sitting in a reclining chair. The experimental session was divided into two parts: 1) wakefulness, in which the chair was set upright and the lights on the room were on, while participants remained awake with their eyes closed and responded to the Simon task for approximately 25 minutes (∼500 trials); 2) drowsiness, in which the chair was reclined, lights were off, and participants were allowed to fall asleep while they kept responding to the experimental task for approximately 1 hour and 30 minutes (∼2000 trials; see Figure 1B). If participants stopped responding for 5 consecutive trials, the experimenter would make a noise to wake them up (e.g., by knocking on the door). The whole session lasted for approximately 3 hours.

Subsequently, for each participant, all trials were classified as “low baseline” or “low arousal” using the algorithmic method developed by Jagannathan and collaborators (2018). The micromeasures algorithm computes the EEG features —predictor variance and coherence— on the pre-trial period (i.e., from -1500 to 0 ms in this particular study), and then classifies trials using a support vector machine (SVM). Predictor variance is assessed across occipital electrodes (O1, Oz, O2) in different frequency bands (2–4Hz; 8–10Hz; 10–12Hz; 2–6Hz), while coherence is measured across specified electrodes in occipital (O1, Oz, O2), frontal (F7, F8, Fz), central (C3, C4), and temporal (T7, T8, TP8, FT10, TP10) regions within delta (1–4Hz), alpha (7–12Hz), sigma (12–16Hz), and gamma (16–30Hz) frequency bands. The SVM uses these features for alertness classification. Specifically, 1) pre-trials containing >50% of alpha oscillations are classified as “awake”, 2) trials containing <50% of alpha oscillations, EEG flattening, and ripples are classified as “drowsy mild”, and 3) further detectors are used to subclassify them into “drowsy severe” (vertex waves, k-complex, and spindles). Finally, to select true awake trials for the “low baseline” condition, we used only trials from the wakefulness blocks and removed all those marked as drowsy (mild or severe). Similarly, drowsy (mild) and drowsy (severe) from the drowsiness blocks were selected as true drowsy trials for the “low arousal” condition.

### Experiment 2

Experiment 2 consisted of three sessions.

#### Incremental effort test

Upon arrival at the testing room, participants signed the informed consent. After a 5 mins warming up at 120 watts, they were asked to pedal on a cycle-ergometer while HR, volume of oxygen (VO_2_) capacity and respiratory exchange ratio were measured. The effort test consisted of increments of 20 watts in power output every 3 minutes, until participants reached full exhaustion and decided to stop. The watts output when the participant reached 80% of their maximum VO2 consumption (VO_2 max_) was used as target watts during the high arousal condition. Whereas, in the baseline condition, the target was 10% of the watts reached at VO_2 max_. Next sessions were held about 1 week after the incremental effort test, to avoid carry-over fatigue effects.

#### Experimental sessions

The two sessions (i.e., high arousal and high baseline) were conducted on two different days, separated by an interval of 24-72 hours (see Figure 1B). The order of both sessions was counterbalanced across participants. Moreover, to control for potential circadian rhythm effects, each individual attended both sessions at the same time of the day.

Upon arrival at the laboratory, participants were fitted with 3 ECG electrodes (Ambu, Ballerup, Denmark) and a 64-channel high-density actiCHamp EEG system (Brain Products GmbH, Munich, Germany). Then, they got on a cycle-ergometer (SRM GmbH, Germany), and were fitted with a HR monitor (SRM GmbH, Germany) and earphones (Kingston, USA). During a 5-minute warm-up, watts were increased every minute or 30 seconds until 80% of VO_2 max_ was reached (high arousal condition) or remained at 10% of VO_2 max_ (high baseline condition). For the first experimental session, during the warm-up, participants were presented a 1-minute practice run of the task (∼20 trials). Next, the experimental task was administered while participants were cycling either at low (10% of VO_2 max_) or high (80% of VO_2 max_) intensity for 30 minutes (∼750 trials per session). Average watts and HR values of the 39 participants for each condition (high arousal and high baseline) are shown in Table 1, to better illustrate the effective intensity of the induced physiological stress and facilitate the comparison of the current findings with existing literature. Individual HR and watts mean values per condition are provided in Supplementary material. The resistance of the bike was controlled by the software MagDays, stimuli were presented using a PsychToolbox software in Matlab, and data were acquired with Brain Vision and AcqKnowledge softwares, and a BIOPAC amplifier, on Dell (USA) computers. Note that EEG data are reported in Avancini et al. (2023).

**Table 1.** Group average HR and watts achieved in each experimental session in Experiment 2, and average percentage relative to their maximum value.

|  | HRmax | VO2max | High arousal |  |  |  |  | High baseline |  |  |  |
| --- | --- | --- | --- | --- | --- | --- | --- | --- | --- | --- | --- |
|  |  |  | Watts at VO2max | Watts | % of max watts | HR | % of HRmax | Watts | % of max watts | HR | % of HRmax |
| Mean | 180.59 | 3.37 | 291.28 | 214.86 | 73.82 | 154.77 | 85.71 | 69.27 | 24.04 | 98.07 | 54.32 |
| 95% CI | [176.3, 184.88] | [3.21, 3.53] | [280.17, 302.39] | [205.1, 224.6] | [71.82, 75.2] | [150.34, 159.21] | [84.34, 87.08] | [64.33, 74.21] | [22.96, 26.02] | [94.75, 101.4] | [52.18, 56.46] |

HRmax: maximum heart rate achieved by participants in the incremental effort test prior to the two experimental sessions. VO2max: maximum volume of oxygen consumption achieved by participants in the incremental effort test.

### Data analyses

#### Reaction times and accuracy

Incorrect or missed trials and trials with RT below 200 ms (i.e., anticipatory responses) were excluded from behavioural analyses^2^. We collapsed data from the two experiments into a single dataset with RT and accuracy as the main indexes of performance. Since the low arousal (i.e., drowsiness and awake) and high arousal (i.e., high- and low-intensity physical exercise) data came from separate databases and different participants, we fitted the variables using multilevel linear mixed-effects modelling, as implemented in the *lme4* R package (Bates et al., 2014). Hierarchical modelling helps minimise any potential differences between the databases that may have not been exclusively caused by the experimental manipulation (e.g., different recording systems or samples) since, unlike traditional analyses, multilevel analysis preserves information about group membership. For instance, it can perform *t*-tests on group-level means, distinguishing between variance within groups (differences among participants within a dataset) and variance between groups (differences among condition means). Thus, multilevel linear mixed-effects modelling effectively addresses the potential similarity of observations from the same experiment by retaining information about each observation’s group membership when assessing parameters such as group differences (Aarts et al., 2014). We treated RT and accuracy as obeying a multilevel data structure, with participant (level 2), nested into experiment (level 1). Participant nested into experiment were set as random factors, and arousal as fixed factor. This structure was common to all models. We performed further models including additional relevant variables (i.e., current congruency, previous congruency) and their interactions to test our hypotheses. All hypotheses were tested using this approach (details on testing model assumptions can be found in the Supplementary material). Additionally, we conducted Bayesian repeated measures t-tests with JASP (version 0.15.0) to examine the favourability towards the null or the alternative hypothesis when splitting per experimental conditions (i.e., comparing high and low arousal conditions with their respective baseline conditions)^3^. For all Bayesian tests, priors were fixed at a Cauchy scale value of 0.4 (i.e., small-to-moderate effect size predicted).

#### Drift-diffusion modelling

DDMs are sequential sampling models which invariably assume that, when an individual faces a two-alternative choice, the amount of evidence for one answer over the other accumulates gradually over time until it reaches a decision threshold, which triggers the motor response (Ratcliff et al., 2016). In this study, we particularly applied a diffusion model for conflict tasks (DMC; Ulrich et al., 2015), which distinguishes between task-irrelevant information or automatic processing (i.e., stimuli location in this scenario) and task-relevant or controlled processing (i.e., word meaning). DMC specifically assumes that, although a single evidence accumulation process combining automatic and controlled processing will determine the executed response, automatic processing will facilitate the controlled accumulation process in congruent trials and will hinder it in incongruent trials. That is, this DDM model presumes that the rate of accumulation (i.e., drift rate) of the information directly related to the task needed to make a decision and provide a response will be affected by a pulse function (which increases and then goes back to zero again) of task-irrelevant processing. For instance, in this particular task, the speed and direction of the accumulation of information about the content of the word (towards a decision threshold: pressing the right or left button) will be affected by the automatic processing of the location at which the word is presented, thereby hindering the accumulation process if the two features are incongruent (please, see Figure 4 for a schematic representation of DMC parameters.

By implementing the *DMCfun* package in R (Mackenzie & Dudschig, 2021), we estimated for each participant under each arousal condition the following parameters: (1) drift rate of controlled processes (i.e., the speed and direction of task-relevant information accumulation), (2) non-decision time accounting for other processes, such as sensory encoding and motor response, (3) distance between decision boundaries (meaning the amount of evidence needed to achieve one of the two decision thresholds, the volume of information used to make the decision regardless of the speed at which it is uptaken), and (4-5) amplitude and time to peak of automatic processing pulse function. Parameters were estimated from fitting the DMC to our data with the differential evolution optimization algorithm (Storn & Price, 1997), relying on the *DEoptim* R package (Mullen et al., 2011). The implemented function *dmcFitSubjectDE* is a global optimization method which fits theoretical data simulated from the model to observed data subject-by-subject, by minimising the root-mean-square error (RMSE) between a weighted combination of the conditional accuracy function (CAF) and the cumulative distribution function (CDF; see Mackenzie & Dudschig, 2021, for further information about the RMSE cost function; see Supplementary material for further details on fitting procedure and parameter recovery tests). As for RT and accuracy data, we analysed the obtained DDM parameters by fitting multilevel linear mixed-effects modelling and Bayesian tests.

## 3. Results

### The impact of arousal on overall task performance

To test whether overall performance was reliably worse in high and low arousal compared to their respective baseline conditions, average RT and accuracy were calculated per participant in each of the experimental conditions. Next, we tested whether arousal influenced overall RT and accuracy. The hierarchical linear mixed-effects model revealed a strong effect of arousal in RT, F (214) = 55.24, *p* < 0.001, ηp^2^ = 0.34, and accuracy, F (235) = 10.24, *p* < 0.001, ηp^2^ = 0.08. When splitting the comparisons to each experiment, reliably slower RT, t (214) = -10.46, *p* < 0.001, *d* = 1.10, BF_10_ = 31166, and lower accuracy, t (239) = 4.50, *p* < 0.001, *d* = 0.79, BF_10_ = 271.61, were only found in low arousal compared to its baseline condition. Neither RT, t (214) = -1.01, *p* = 0.32, *d* = 0.12, BF_10_ = 0.35, nor accuracy, t < 1, BF_10_ = 0.298, showed reliable differences between high arousal and its baseline (see Figure 2A-D). Hence, overall performance was negatively impacted only at the lower side of the arousal spectrum.

**Figure 2:**
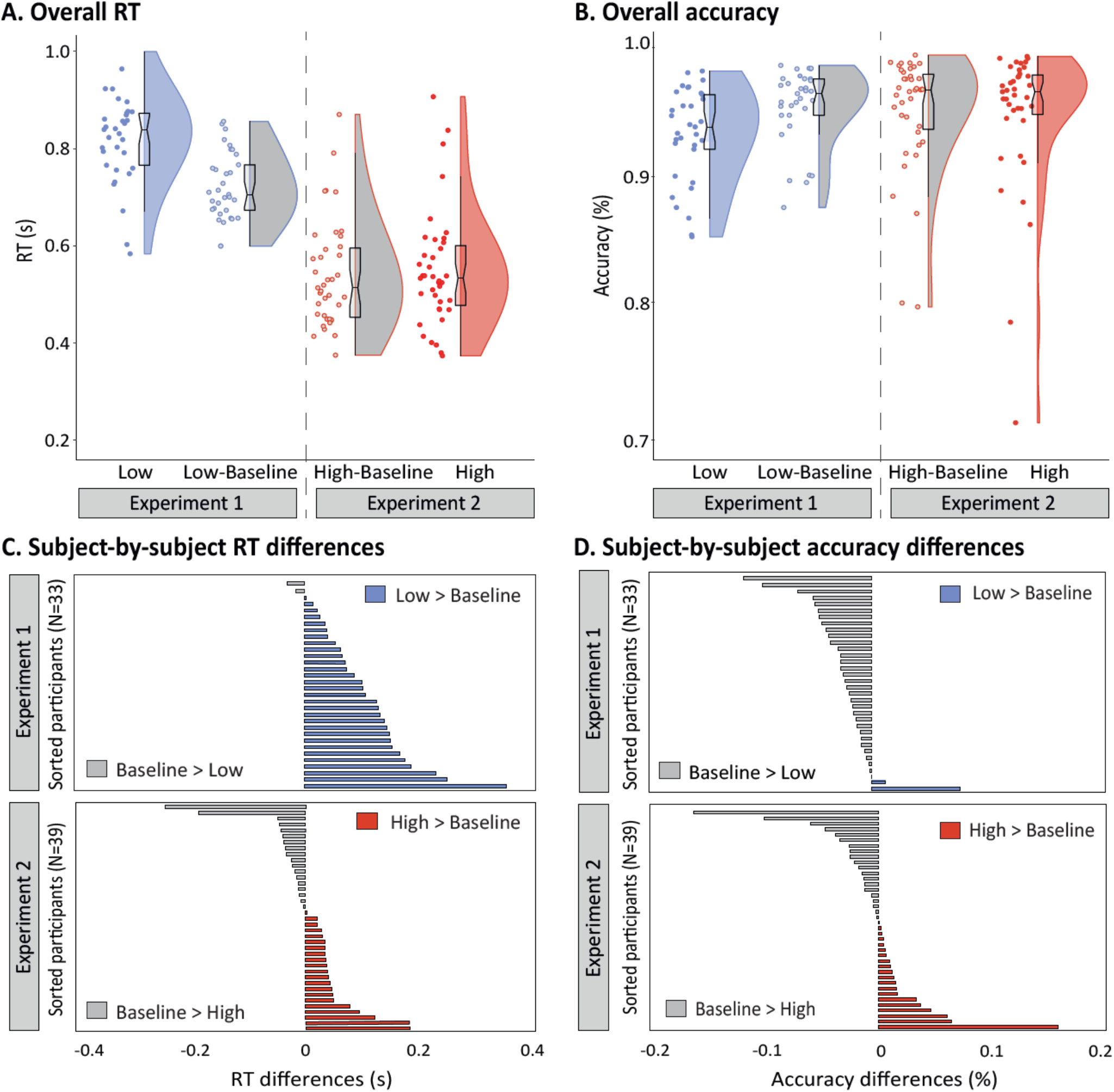
Overall RT and accuracy main results. A-B) Violins and overlaid box plots of mean overall RT (A) and accuracy (B) across the arousal conditions (low arousal and its baseline in Experiment 1, and high arousal and its baseline in Experiment 2). Accuracy is represented as a ratio between 0 and 1, where 1 indicates 100% correctness (i.e., no errors). In box plots, the middle black mark indicates the median, and bottom and top edges indicate 25th and 75th percentiles, respectively. The upper and lower whiskers indicate the distance of 1.5 times the interquartile range above the 75th percentile and below the 25th percentile. Adjacent to the box, on the right, there is a rotated kernel density plot; on the left, jittered dots depict the average RT (A) or accuracy (B) of each individual in each arousal state. C-D) Global RT (C) and accuracy (D) differences of each participant in both altered arousal states (i.e., high and low arousal), compared to their respective baselines. Participants are sorted by performance difference between baseline and the arousal state (i.e., the outcome of subtracting the value of RT/ accuracy in the respective baseline condition from its value in the high or low arousal condition). Grey bars indicate that these participants showed higher RT/accuracy in the baseline condition, compared with the altered arousal states; while blue and red bars depict that these participants showed slower RT or higher accuracy in the low or high arousal states, compared with their respective baselines.

### Cognitive control preservation under altered arousal states

In order to test whether arousal had an impact on the conflict effect, the average RT and accuracy per participant for congruent and incongruent trials were calculated and fitted using a hierarchical linear mixed-effects model, with arousal, congruency and its interaction (i.e., ‘arousal x congruency’) as fixed effects. As predicted, RT yielded a robust effect of congruency, F (211) = 46.06, *p* < 0.001, ηp^2^ = 0.18, but no reliable interaction between arousal level and current trial congruency, F (211) < 1, *p* = 0.58, ηp^2^ = 0.005 (see Figure 3A). To further examine the favourability towards the absence of differences in conflict effect between arousal conditions, we split the data per arousal conditions and performed additional frequentists and Bayesian tests. No reliable differences were found in the magnitude of congruency effect on RT between high arousal and its baseline, t (211) < 1, *p* = 0.54, *d* = 0.24, BF_10_ = 0.69, as well as between low arousal and its baseline, t (211) < 1, *p* = 0.55, *d* = 0.21, BF_10_ = 0.53. Thus, although Bayesian contrasts yielded inconclusive evidence, global results point towards a relative preservation of the RT conflict effect in both high and low arousal states compared to their baseline conditions (see Figure 3C). Accuracy analysis revealed a reliable main effect of congruency, F (211) = 51.21, *p* < 0.001, ηp^2^ = 0.15, but no reliable interaction between arousal and congruency, F (211) = 3.07, *p* = 0.49, ηp^2^ = 0.03. When comparing high and low arousal conditions with their respectives baselines, no differences were found in conflict effect between low arousal and its baseline, t (211) = 1.59, *p* = 0.11, *d* = 0.16, BF_10_ = 0.43, or between high arousal and its baseline, t (211) = -1.27, *p* = 0.21, *d* = 0.06, BF_10_ = 0.30.

**Figure 3:**
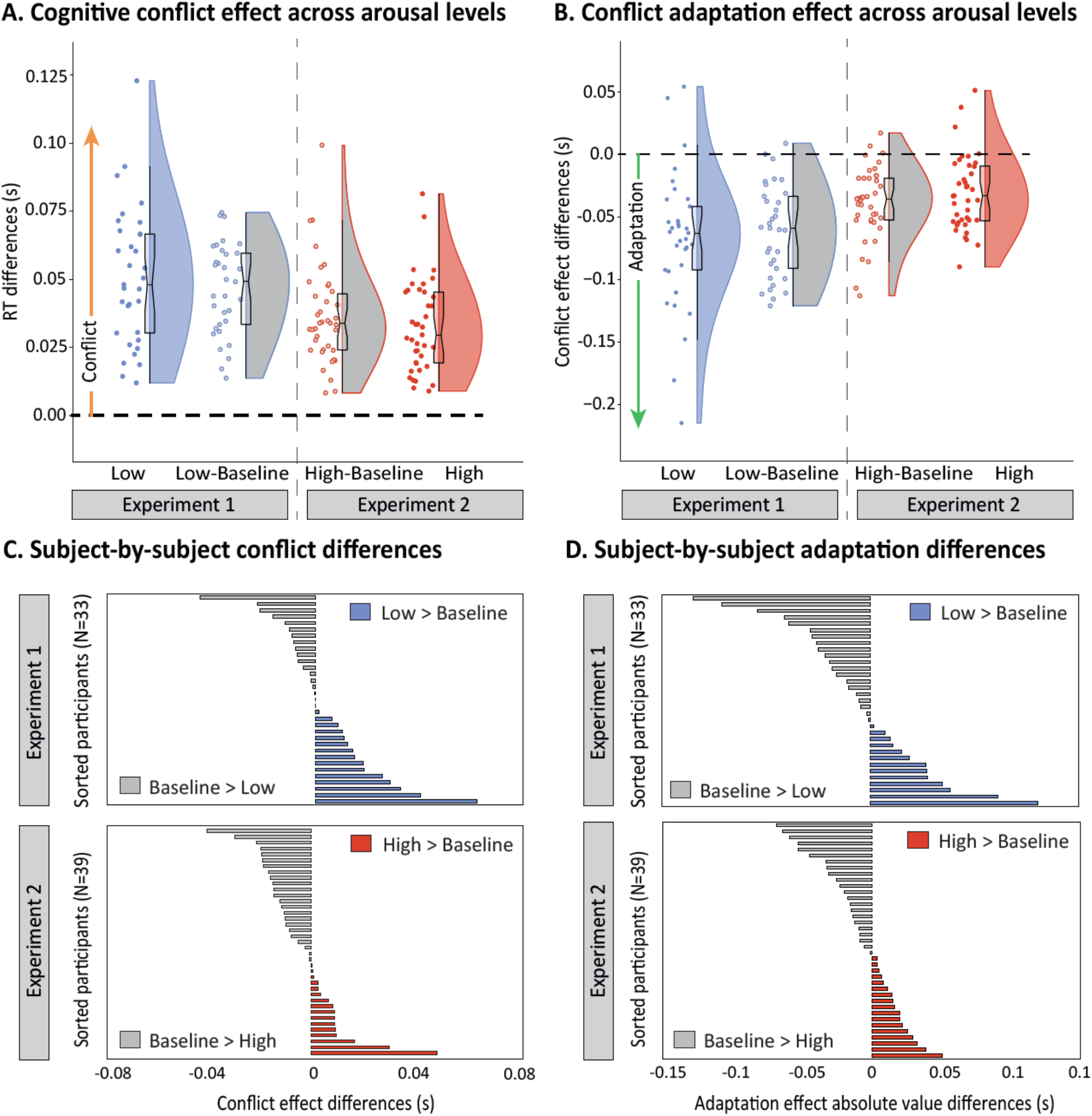
Current and previous trial congruency effects as a function of arousal level. A) Violins and overlaid box plots of RT conflict effect (i.e., RT for incongruent trials minus RT for congruent trials) for each arousal condition (low arousal and its baseline in Experiment 1, and high arousal and its baseline in Experiment 2). In box plots, middle black mark indicates the median, and bottom and top edges indicate 25th and 75th percentiles, respectively. The upper and lower whiskers indicate the distance of 1.5 times the interquartile range above the 75th percentile and below the 25th percentile. Adjacent to the box, on the right, there is a rotated kernel density plot; on the left, jittered dots represent the average RT conflict effect of each individual in each arousal state. Every participant showed a global conflict effect in all experimental conditions and, as expected, no statistically reliable differences were found in the magnitude of the conflict effect between the altered arousal states and their respective baseline conditions. B) Violins and overlaid box plots of RT conflict adaptation effect (i.e., conflict effect for previous incongruent trials minus conflict effect for previous congruent trials). According to linear mixed-effects analyses, conflict adaptation is preserved in every arousal condition, and there are no statistically reliable differences in the magnitude of conflict adaptation effect between high arousal and high baseline, or low arousal and low baseline. C-D) Global conflict (C) and adaptation absolute values (D) differences of each participant in both altered arousal states (i.e., high and low arousal), compared to their respective baselines. Participants are sorted by performance difference between baseline and the arousal state (i.e., the value of the conflict/adaptation effect in the high or low arousal condition minus its value in the respective baseline condition). Grey bars indicate that these participants showed higher conflict/adaptation in the baseline condition, compared with the altered arousal states. Blue and red bars depict that these participants showed higher conflict/adaptation effect in the low or high arousal states, compared with their respective baselines.

**Figure 4:**
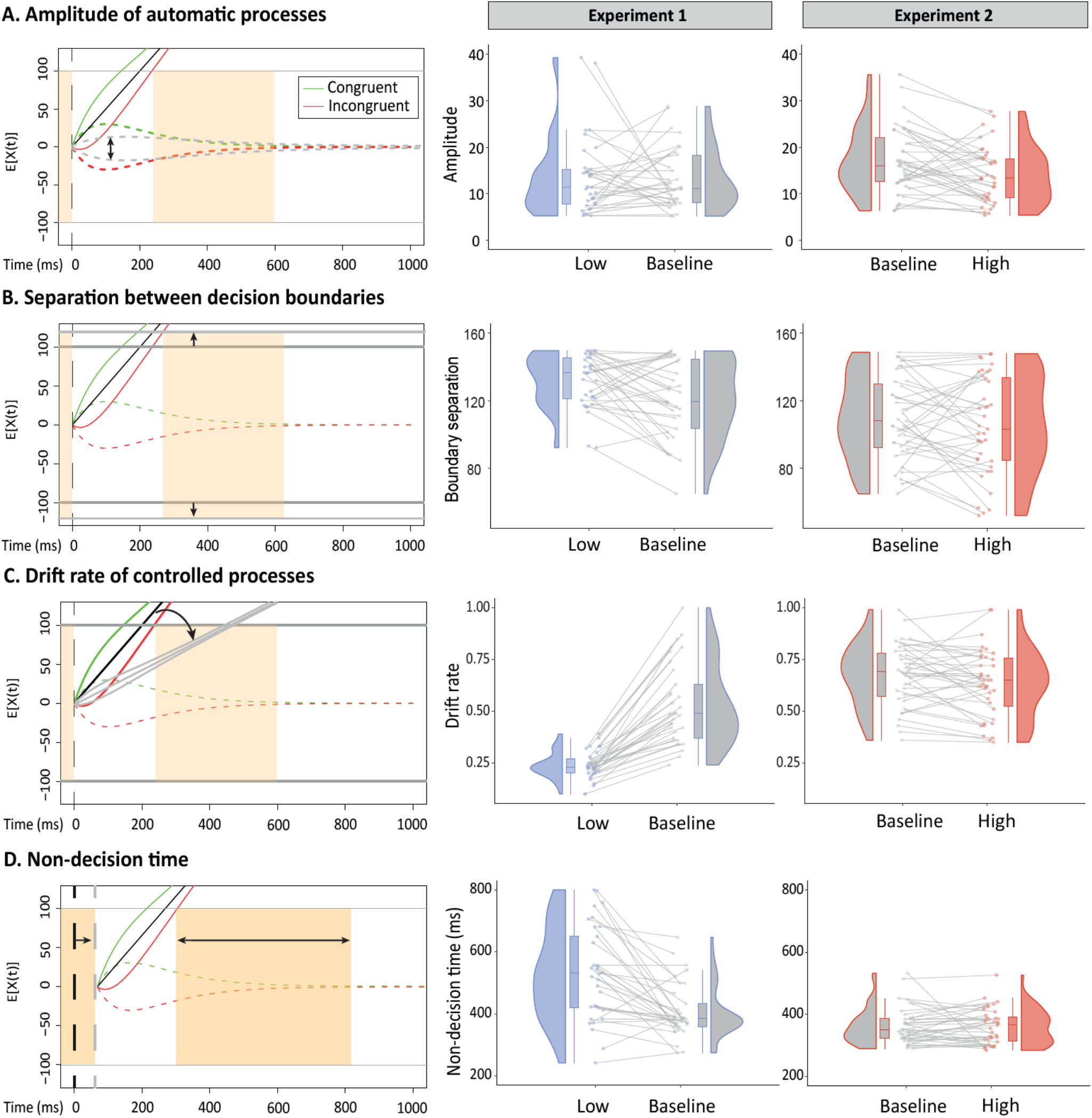
The impact of arousal on decision-making parameters. Panels on the left show the schematic representation of a shift in different diffusion model for conflict tasks (DMC) parameters: (A) amplitude of automatic (i.e., task-irrelevant) processing, (B) boundary separation, (C) rate of evidence accumulation of the controlled (i.e., task-relevant) processing, and (D) non-decision time (i.e., sensory encoding and motor response duration). The DMC illustrative representation on which the changes in the respective parameters have been plotted was generated from the simulation of 100,000 congruent and incongruent trials. Upper and lower grey solid lines represent the boundaries of the two potential responses (correct, above, and incorrect, below), and orange areas depict non-decision time. The black solid line constitutes the drift rate of controlled processing, while the dotted green and red lines represent the automatic processing for congruent and incongruent trials, respectively. The response will be determined by a single rate of evidence accumulation, resulting from the combination of the automatic and controlled processes, represented by the green solid line for congruent trials, and the red one for incongruent trials. Panels in the middle (results from Experiment 1) and on the right (results from Experiment 2) represent subject-by-subject baseline differences for each DMC parameter. Blue, grey and red areas are rotated kernel density plots (i.e., a histogram with infinitely small bin sizes). In box plots, middle marks indicate the median, and bottom and top edges indicate 25th and 75th percentiles, respectively. The upper and lower whiskers indicate the distance of 1.5 times the interquartile range above the 75th percentile and below the 25th percentile. Jittered dots represent the average DMC parameter value (e.g., amplitude of automatic processes in A) of each individual under each arousal state. The two values from the same subject in both experimental conditions are linked by a solid grey line. Linear mixed-effects and Bayesian analyses showed that the amplitude of automatic processes was reliably reduced during high arousal compared to its respective baseline condition, while low arousal was solidly associated with (1) slower drift rate, (2) wider separation of decision boundaries and (3) longer non-decisional time than its baseline.

Similarly, the impact of arousal on conflict adaptation was assessed by conducting a hierarchical mixed-model fitting RT, with arousal, congruency, previous congruency and all their interactions as fixed factors. As hypothesised, the model showed a robust effect of the interaction between current trial congruency and previous trial congruency, F (493) = 45.10, *p* < 0.001, ηp^2^ = 0.08. This interaction actually reflects that, RT for congruent trials were shorter when preceded by congruent trials compared to incongruent ones, and inversely (RT were shorter for incongruent trials preceded by an incongruent trial), which results in a decreased conflict effect when the previous trial was incongruent (i.e., conflict adaptation). Again, hierarchical linear mixed-effects model revealed no reliable effect for the ‘arousal x congruency x previous congruency’ interaction, F (493) = 1.38, *p* = 0.25, ηp^2^ = 0.005 (see Figure 3B). Consistently, the comparison between altered arousal states and their baselines also yielded no reliable differences in conflict adaptation effect, t <1, *p* = 0.6, *d* = 0.29, BF_10_ = 0.89 (low arousal vs. baseline), t (493) = -1.31, *p* = 0.19, *d* = 0.26, BF_10_ = 0.79 (high arousal vs. baseline; see Figure 3D).

### High and low arousal modulate decision-making parameters differently

To further assess the influence of arousal level on decision-making processing, we fitted a DDM for conflict tasks (see section 2 “Data analyses. Drift-diffusion modelling” for further information on model fitting and assumptions) to our databases (see Table 2). We estimated for each participant under each arousal condition five parameters: (1) drift rate of controlled processes (i.e., the speed and direction of task-relevant information accumulation), (2) non-decision time (sensory encoding and motor response), (3) distance between decision boundaries (i.e., amount of evidence needed to achieve one of the two decision thresholds), and (4-5) amplitude and time to peak of automatic (task-irrelevant information) processing pulse function. The hierarchical linear mixed-effects model revealed a robust effect of arousal level in drift rate of controlled processes, F (70) = 70.9, *p* < 0.001, ηp^2^ = 0.67, separation between decision boundaries, F (62) = 5.28, *p* = 0.008, ηp^2^ = 0.14, and non-decisional time, F (59) = 24.22, *p* < 0.001, ηp^2^ = 0.45. The effect of arousal on automatic processing parameters was not statistically reliable, F (63) = 2.27, *p* = 0.11, ηp^2^ = 0.07 (amplitude), F (91) = 1.71, *p* = 0.19, ηp^2^ = 0.04 (time to peak).

**Table 2.** Group average of DMC estimated parameters across the arousal conditions.

|  | Experiment 1 |  |  |  | Experiment 2 |  |  |  |
| --- | --- | --- | --- | --- | --- | --- | --- | --- |
|  | Low arousal |  | Low baseline |  | High baseline |  | High arousal |  |
|  | Mean | 95% CI | Mean | 95% CI | Mean | 95% CI | Mean | 95% CI |
| **Amplitude (automatic processes)** | 13.62 | [10.64, 16.6] | 13.19 | [10.87, 15.5] | 17.12 | [14.86, 19.39] | 14.05 | [12.1, 16] |
| **Duration (automatic processes)** | 151.53 | [112.2, 190.85] | 129.99 | [98.89, 161.08] | 143.17 | [114.6, 171.75] | 167.66 | [138.63, 196.69] |
| **Drift rate (controlled processes)** | 0.24 | [0.22, 0.26] | 0.53 | [0.46, 0.6] | 0.67 | [0.63, 0.72] | 0.63 | [0.58, 0.69] |
| **Boundary separation** | 131.92 | [126.39, 137.45] | 120.79 | [112.62, 128.97] | 110.16 | [102.23, 118.1] | 106.12 | [96.67, 115.57] |
| **Non-decision time** | 529.34 | [476.39, 582.28] | 407.83 | [376.79, 438.86] | 362.77 | [344.04, 381.49] | 361.14 | [342.82, 379.46] |
| **Fitting cost value (RMSE)** | 84.18 | [74.73, 93.61] | 25.08 | [18.94, 34.78] | 10.47 | [6.99, 14.1] | 14.07 | [8.67, 19.46] |

95% CI: 95% confidence interval of mean. RMSE: root-mean-square error.

Following the same rationale as in previous analyses, Bayesian comparisons between arousal conditions and their baselines were undertaken to examine the weight of evidence supporting the absence/presence of reliable differences in decision-making parameters across the arousal spectrum. Splitting per arousal conditions, mixed-effects modelling showed a statistically significant difference in the amplitude of automatic processes between high arousal and its baseline condition, t (69) = 2.12, *p* = 0.04, *d* = 0.49, a result that was further supported by Bayesian analysis that showed evidence for the alternative, BF_10_ = 9.41. Based on these results, the amount of interference from non-relevant information would be slightly reduced during high arousal state, compared to its baseline (see Figure 4A). While most Bayesian analyses yielded inconclusive evidence, frequentist statistics did not find reliable differences for the rest of the DMC parameters: duration of automatic processes, t (90) = -1.76, p = 0.08, *d* = 0.29, BF_10_ = 1.12; drift rate of controlled processes, t (70) = 1.70, p = 0.09, *d* = 0.32, BF_10_ = 1.4; boundary separation, t (69) = 1.26, p = 0.21, *d* = 0.16, BF_10_ = 0.431; and non-decision time, t (68) < 1, *p* = 0.77, *d* = 0.11, BF_10_ = 0.29.

When compared with its baseline, low arousal was associated with a reliably lower drift rate of controlled processes, t (70) = 11.797, p < 0.001, *d* = 1.815, BF_10_ = 6.9e+8, a wider separation of decision boundaries, t (67) = -2.95, p = 0.004, *d* = 0.50, BF_10_ = 6.62, and a higher non-decisional time, t (67) = - 6.94, p < 0.001, *d* = 0.85, BF_10_ = 641.73. Automatic processing parameters did not yield statistically reliable differences between low arousal and low baseline, t (68) < 1, *p* = 0.92, *d* = 0.06, BF_10_ = 0.39 (amplitude), t (95) < 1, *p* = 0.35, *d* = 0.15, BF_10_ = 0.42 (time to peak). These results suggest that, during drowsiness, evidence accumulation and non-decision time would become slower, while more evidence might need to be accumulated to reach a response (see Figure 4B-D).

## 4. Discussion

In the present study, healthy participants performed a “conflict task” while transitioning into sleep (drowsiness) or physical extenuation (high-intensity physical exercise) to study the behavioural dynamics of cognitive control during non-pharmacological altered arousal states. In line with our pre-registered hypothesis, conflict and conflict adaptation effects were preserved during high and low arousal. While overall task performance was poorer at the lower side of the spectrum (i.e., in the low arousal condition compared to its baseline), this impairment was not observed at the upper side (i.e., high arousal condition). DDM analyses revealed that the impairment of performance during low arousal might be due to: (1) a slower rate of task-related information evidence accumulation; (2) a longer non-decision time that could be caused by a deceleration in sensory encoding, motor response execution, or both; and (3) a wider separation between decision thresholds, which means that more evidence would need to be accumulated by the system to reach a response criterion. Notably, although increased levels of arousal showed minimal performance changes, they were associated with a decrease in the amplitude (i.e., interference) of task-irrelevant information processing. In general, these results provide evidence that high and low arousal differentially impact overall performance and the underlying decision-making processes during automated tasks and reveal how cognitive control-related mechanisms change.

It has been previously shown how decreased arousal fails to fully interrupt higher order cognitive processes, such as perceptual decision-making (Bareham et al., 2014, 2015), semantic discrimination (Kouider et al., 2014) or probabilistic learning (Ciria et al., 2021). However, as reported by the entire body of literature on drowsy states (Bareham et al., 2015; Canales-Johnson et al., 2020; Ciria et al., 2021; Goupil & Bekinschtein, 2012; Jagannathan et al., 2022; Kouider et al., 2014; Noreika et al., 2020; Xu et al., 2023), decreasing the level of arousal leads to slower RT, lower accuracy and decreased sensitivity in decision-making. Our results replicate and enable the generalisation of previous reports, further revealing that drowsiness selectively modulates task-relevant decision-making processes but fully preserves some automated aspects of cognitive control. Notably, EEG analyses of this same database (Canales-Johnson et al., 2020) show that the robust behavioural and computational effects of efficient conflict resolution are not accompanied by their classical neural marker (i.e., increased power in mid-frontal theta), and suggest some reorganisation of the traditional (waking) neural activity to deal with conflict during a non-homeostatic state (i.e, drowsiness).

The transition towards the other side of the arousal spectrum, however, yielded mixed results. Consistent with previous studies examining cognitive control during moderate-to-high physical exercise (Davranche et al., 2009, 2015; Joyce et al., 2014; Schmit et al., 2015), conflict detection and adaptation were both fully preserved during high arousal. As in low arousal, this was not accompanied by the preservation of the classical neural correlate of conflict resolution (Avancini et al., 2023), suggesting some parallelisms between these two strained states of arousal. Yet, the absence of impaired cognitive performance when participants exercised close to the anaerobic threshold deviates from the hypotheses initially proposed. Some accounts have shown that acute exercise might not produce a deterioration in performance when the task involves highly automated behaviours or rapid decision-making (Lambourne & Tomporowski, 2010). The Simon task may meet both requirements, as participants had to respond as quickly as possible to two highly automated words. From a theoretical perspective, these findings would therefore be in line with the task-difficulty assumption within the Yerkes-Dodson framework, which proposes easier tasks require higher levels of arousal for optimal performance than more difficult tasks (Sjöberg, 1977). However, it is important to note that the Yerkes-Dodson proposal is utilised solely as a useful starting point for studying the relationship between arousal and cognition. Initially, this law was not formulated for this purpose but rather in the context of learning. The generalisation from its original formulation (Teigen, 1994) and its reductionist nature may pose a limitation in the interpretation of results from this perspective. Furthermore, the current state of the literature on physical activity, containing either studies in which cognitive performance does worsen during moderate-to-high exercise (Chmura et al., 1997; Ciria et al., 2021; Del Giorno et al., 2010; González Fernández et al., 2017; Zheng et al., 2021) or studies where performance remains the same or even improves (e.g., Lambourne & Tomporowski, 2010; Sudo et al., 2022), raise the question of whether this theory, and even the wider notion of arousal, is sufficiently informative when interpreting performance outcomes.

As opposed to low arousal, high arousal was found to have no impact on rate of evidence accumulation, boundary separation and non-decision time. Instead, our analyses revealed unexpected differences in the amplitude of automatic processes between high arousal and its baseline condition, whereby the amount of interference from task-irrelevant (conflicting and non-conflicting) information would be reduced during this state. The scarce literature on how decision parameters are modulated by high arousal makes it difficult to assess the reliability of these findings. To our knowledge, only two studies have addressed decision-making during physical exercise using a DDM approach (Lefferts et al., 2016; 2019). In these studies, participants performed a flanker task while (or immediately after) exercising at moderate intensity. The authors reported no reliable differences on any of the estimated cognitive parameters. Notably, the DDM in their research did not distinguish between automatic and controlled processing, and therefore it could not detect the effect in the amplitude of automatic processes reported here, even if it was present. Interestingly, EEG multivariate spectral decoding analyses of the high arousal data showed that the task-irrelevant information (i.e., spatial location) was only present (decodable) in the baseline condition in the 2-10 Hz band, but not during high-intensity exercise (Avancini et al., 2023). Also consistent with a narrower focus during strained states of high arousal, reduced overt attentional capture (i.e., automatic disruption from abrupt distracting stimuli) has been previously observed right after a bout of intense physical exercise (Llorens et al., 2015). Thus, although further research is needed to draw more solid conclusions, increased arousal seems to produce little difference in classical behavioural indices, whereas computational and brain activity outcomes would provide a hint that the system undergoes some cognitive modulations during such state, albeit in a different direction to that of low arousal.

The present study highlights the utility of inducing natural states of drowsiness and physical exercise as causal models to study cognition as our level of arousal moves towards either of its extremes. Despite their different biological nature, both transitions —towards sleep and physical exertion— involve rapid and dynamic fluctuations in arousal, accompanied by important physiological and phenomenological changes, in which the overall capacity to respond behaviourally to stimuli is preserved (Ciria et al., 2021). These features, together with the implementation of multilevel mixed-effects models, allow us to combine them in a common experimental framework to study how human cognitive and neural dynamics are modulated as our system becomes overstrained on a physiological level. Note, however, that although hierarchical modelling attempts to minimise potential differences between databases (i.e., experiments) when estimating overall arousal effects, drowsiness and physical exertion are not directly compared to each other in any part of the study. Future studies in which a single sample experiences all physiological states could result in more conclusive findings. On the other hand, there is a growing consensus in neuroscience that RT and accuracy, while undoubtedly useful, can sometimes be insufficiently informative about performance (Voss et al., 2013). Indeed, our findings showcase the benefits of using computational modelling as a tool to identify the specific decision-making mechanisms underlying the modulations observed through classical measures (e.g., RT). Future steps involve further deepening the complementary nature of neural and computational analyses to disentangle the cognitive dynamics during everyday arousal fluctuations. For now, the combination of drowsiness and intense physical exercise, together with the use of behavioural computational modelling, seem to offer a fruitful approach to explore the cognitive processing during endogenously altered arousal states.

## Supporting information

Supplementary_material_Alameda_2024

## 5. Acknowledgements and funding

This research was supported by a predoctoral fellowship by the Spanish Ministry of Universities to the first author (FPU21/00388), a postdoctoral fellowship by the Spanish Ministry for Science and Innovation to the second author (FJC2020-046310-I); a research project grant from the Spanish Ministry of Science and Innovation to D.S. (PID2019-105635GB-I00), a postdoctoral fellowship from the Regional Government of Andalusia to L.F.C. (DOC_00225), and a Wellcome Trust Biomedical Research Fellowship (WT093811MA) awarded to T.A.B. We thank Luc Vermeylen for his valuable help with the drift-diffusion modelling analyses.

## 6. Open Practices Statement

The hypotheses and analyses plan were pre-registered in AsPredicted after data collection but prior to data observation and analysis (https://aspredicted.org/x8qd6.pdf), except for DDM analyses. All deviations from the pre-registered procedures are transparently identified in the manuscript. Raw data, sorted data and codes used for the analyses presented here are available at the OSF repository (https://osf.io/uvq3m/).

## 7. CRediT Author Statement

**Clara Alameda:** Conceptualization, Methodology, Software, Data curation, Formal Analysis, Visualization, Writing, Reviewing and Editing. **Chiara Avancini:** Conceptualization, Methodology, Data curation, Software and Reviewing. **Daniel Sanabria:** Conceptualization, Methodology, Reviewing, Editing, and Supervision. **Tristan A. Bekinschtein:** Conceptualization, Methodology, Visualization, and Reviewing. **Andrés Canales-Johnson:** Conceptualization, Methodology, and Reviewing. **Luis F. Ciria:** Conceptualization, Methodology, Software, Visualization,, Reviewing, Editing and Supervision.

## Footnotes

1 In Experiment 1, trials were labelled according to three dimensions, being wakefulness (awake vs. drowsy), current trial congruency and previous trial congruency (congruent vs. incongruent), thereby creating eight conditions. After labelling, 8 participants from Experiment 1 were excluded from subsequent analyses because of insufficient data (i.e., less than 10 trials in a single condition).

2 Data exclusion procedure not included in the pre-registration. Following Canales-Johnson and collaborators’ (2020) behavioural data processing, we excluded anticipatory responses, errors and omissions.

3 Bayesian analyses were not included in the pre-registration. The analyses were conducted after multilevel models aiming to test those hypotheses concerning the absence of differences.

