## Supplementary_material_Alameda_2024 for "Staying in control: characterising the mechanisms underlying cognitive control in high and low arousal states"

**DDM fitting procedure and reliability**

***Fitting real data with a differential evolution algorithm***

Decision-making parameters for each subject under each arousal condition were estimated from fitting the diffusion model for conflict tasks (DMC; Ulrich et al., 2015) to our data using the differential evolution optimization algorithm of Storn and Price (1997) within the R packages *DMCfun* (Mackenzie & Dudschig, 2021) and *DEoptim* (Mullen et al., 2011). The implemented function *dmcFitSubjectDE* is a global optimization method which fits theoretical data simulated from the model to observed data subject-by-subject, by minimising the root-mean-square error (RMSE) between a weighted combination of the conditional accuracy function (CAF) and the cumulative distribution function (CDF; see Mackenzie & Dudschig, 2021, for further information about the RMSE cost function).

We fitted 5 parameters (amplitude of automatic processes, time to peak of automatic processes, drift rate of the controlled processes, decision boundary where the response is executed, and mean of the normal distribution of the non-decision time), which we intended to study along the arousal spectrum, as well as the standard deviation of the non-decision time distribution, allowing to have some inter-trial variability in non-decision time. The remaining 3 parameters of the model (i.e., shape parameter of the gamma distribution for time to peak of automatic processes, shape parameter of the beta distribution for starting point variability, and scaling parameter of the drift diffusion process) were fixed at the *DMCfun* default value, to increase the focus on the estimation of the most informative parameters within the model. Parameters default minimum and maximum values within *DMCfun* were not modified, except for the amplitude of the automatic processes (set at 5-40) and the standard deviation of the non-decisional time (set at 40-200), due to the large variability and the extended RT observed in low arousal data (see Mackenzie & Dudschig, 2021, for further information about *DMCfun* default values). The number of iterations was set at 250. This fitting procedure was equally performed on the data from each of the arousal conditions (i.e., high arousal, high baseline, low baseline and low arousal). Additionally, although the differences between observed and predicted RT and error distributions did not appear to be substantial (see Supplementary figures 1-2), a parameter recovery test with the same fitting settings was performed to ensure the reliability of the estimated parameters (see Parameter recovery below).

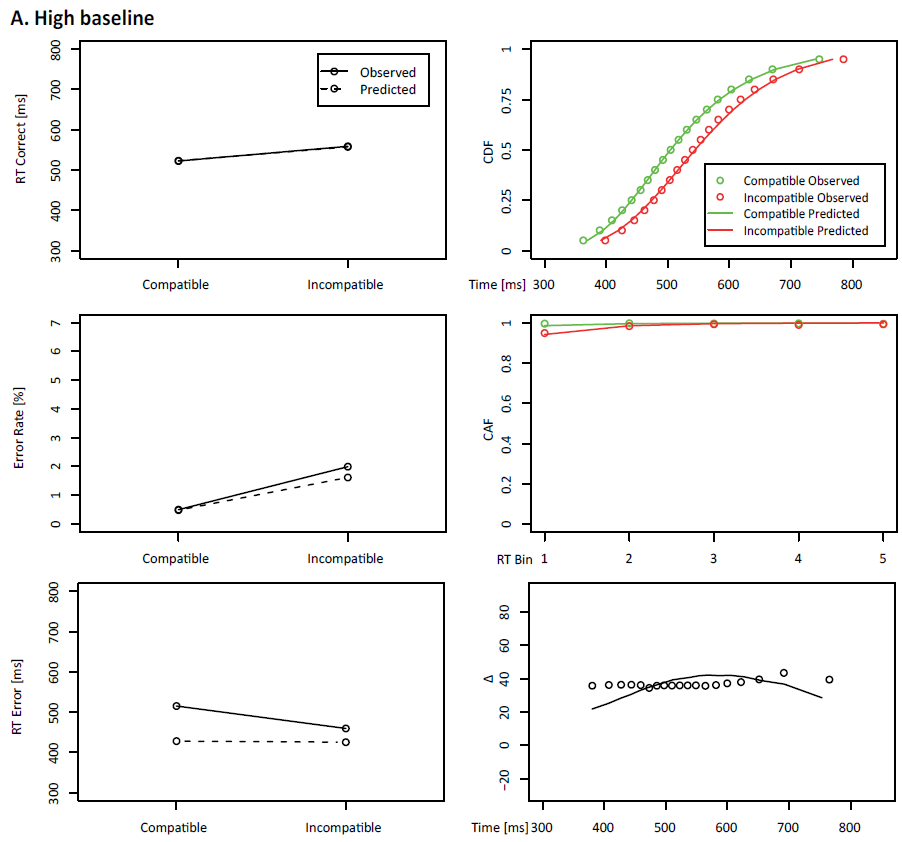

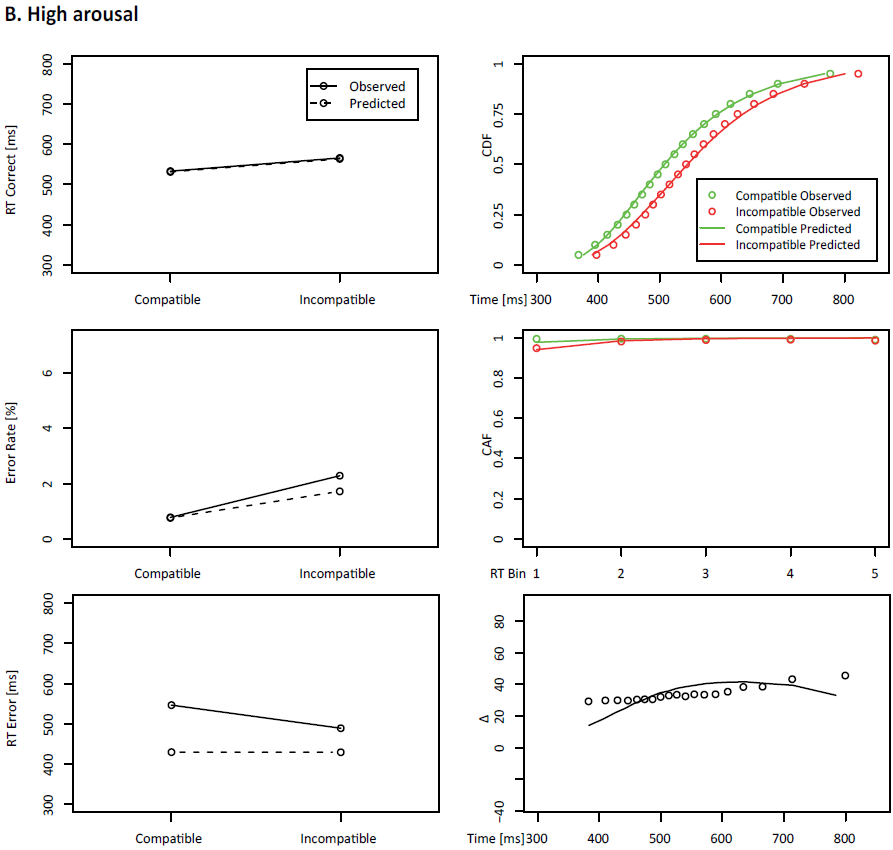

**Supplementary figure 1. DMC fits from the (A) high baseline and (B) high arousal data.** For each condition, the panels along the left-hand column show the observed group mean from the data (linked by solid lines) and the predicted group mean from the model fit (linked by dotted lines) for correct trials RT, error rate, and errors RT in congruent and incongruent trials. The panels on the right column show the RT cumulative distribution function (CDF), the conditional accuracy function (CAF) and the delta plot for the observed (hollow dots) and predicted (solid lines) congruent (green) and incongruent (red) trials. Delta plots represent the average conflict effect as a function of RT. It should be noted that, while the fitting procedure was conducted on a subject-by-subject basis, it is depicted here at the group level. In line with the low fitting cost values (i.e., RMSE), these plots would globally suggest a good model fit to our data.

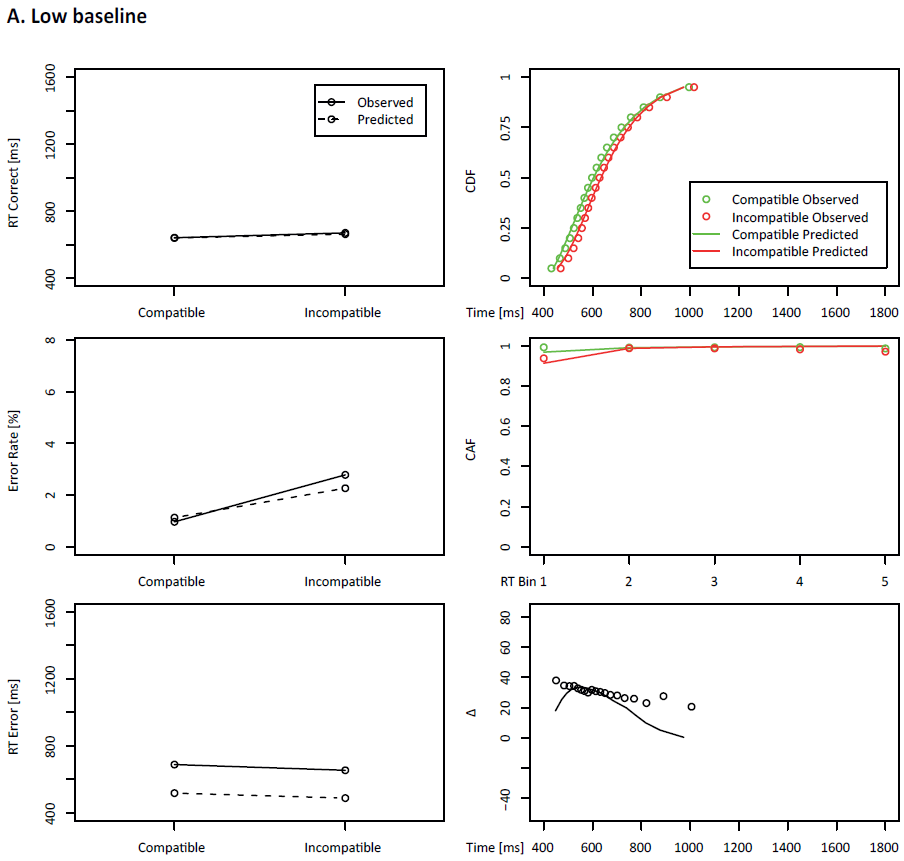

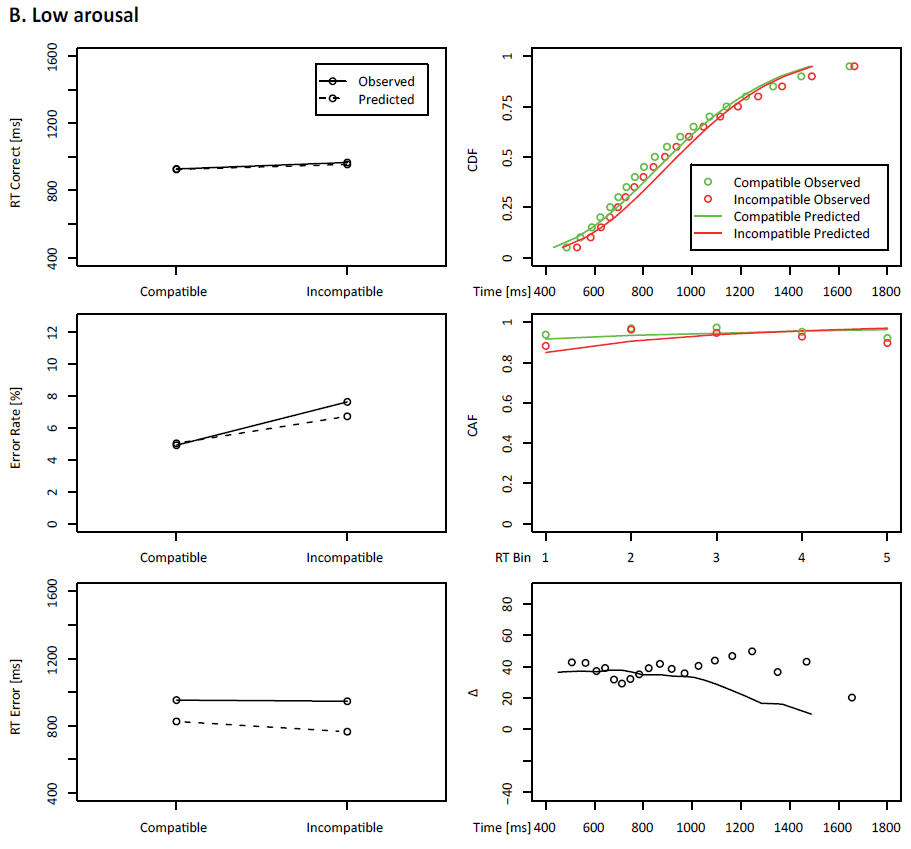

**Supplementary figure 2. DMC fits from the (A) low baseline and (B) low arousal data.** These panels would broadly suggest a reasonable model fit to our low arousal and low baseline data.

***Parameter recovery test***

To examine the accuracy of the DMC parameter estimates, we conducted a parameter recovery test. First, a random set of DMC parameters were drawn from a uniform distribution within the boundaries of the parameter values obtained in our subject-by-subject fitting results. We then simulated a 800-trials dataset (comprising 400 congruent and 400 incongruent trials, similar to the average number of trials per subject in our data), which conforms to the generated parameters. Next, DMC was fitted to the simulated data, maintaining the exact same fitting procedure described above (see Fitting real data with a differential evolution algorithm). The entire process was repeated 100 times. The initial generated parameters (i.e., true values) and the latter parameters resulting from the fitting (i.e., recovered values) were subsequently correlated to see if the implemented fitting procedure successfully retrieved the true parameter values. The results point towards a reliable recovery of the true values for the five DMC parameters examined in this study (see Supplementary figure 3).

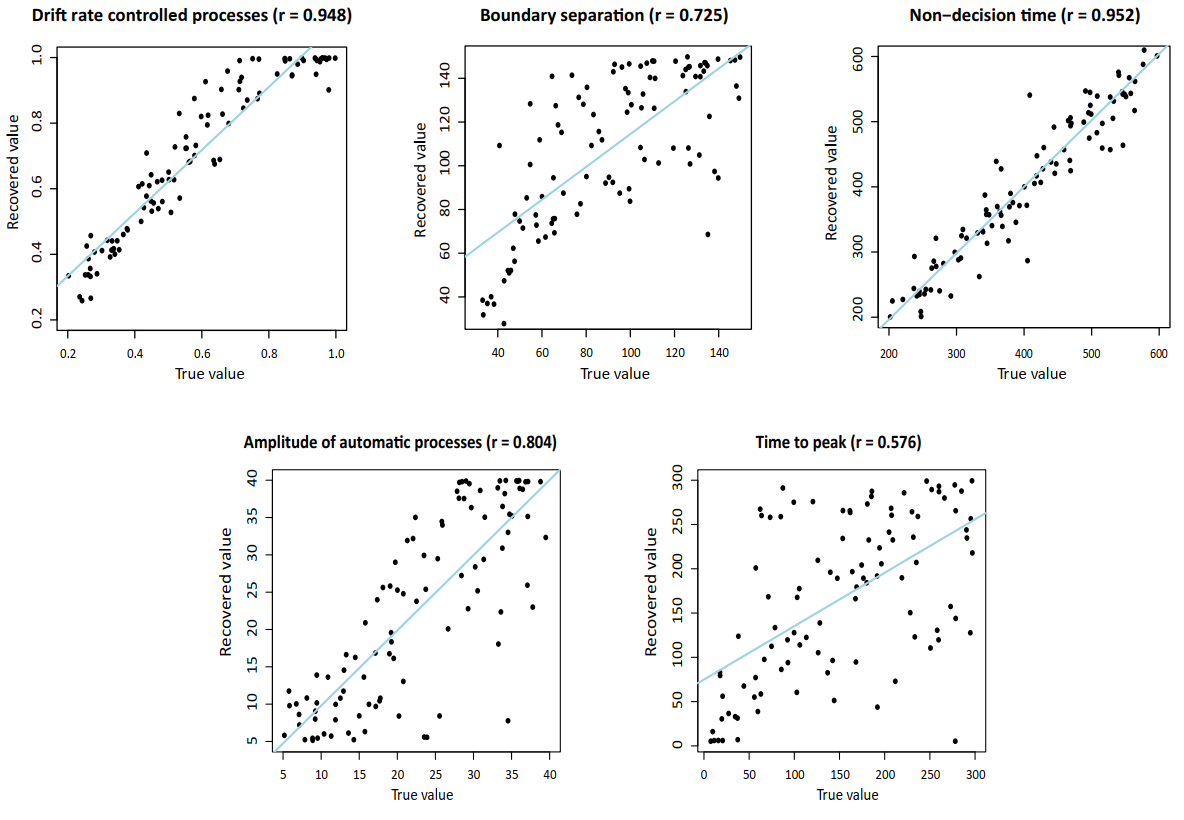
**Supplementary figure 3. Parameter recovery for DMC parameters.** We simulated 100 datasets containing 800 trials (50% congruent and 50% incongruent) from a set of random parameters that fell within the ranges of the parameter values obtained in our results. Next, we fitted the model to the simulated data with the exact same procedure as for the experimental data, to examine the reliability of the DMC parameter estimates. The five panels show the Pearson correlations between the generated and the recovered parameter values for the drift rate of controlled processes, the separation between decision boundaries, the non-decision time (above), and the amplitude and time to peak of the automatic processes (below).

**Testing the assumptions of the generated mixed-models**

***Models for testing RT and accuracy hypotheses***

We tested the main assumptions of mixed models on the two hierarchical linear mixed-effects models we used to examine the effect of arousal (as the only fixed effect) on overall RT and accuracy (herein referred to as ‘ACC’), with participant nested into dataset as random effects. We first tested the linearity of the data by plotting the RT and ACC models’ residuals (i.e., the difference between the observed value and the model-estimated value) versus the predictors (see Supplementary figure 4A-B). The relationship between overall RT and arousal (Shapiro-Wilk normality test = 0.988, p = 0.021) and the relationship between overall ACC and arousal (W = 0.942, p < 0.001) could not be described by a straight line, but by a quadratic function as noted in the Results section. Next, as regression models assume that variance of the residuals is equal across groups, we checked whether the variance of residuals was equal across datasets (i.e., Experiment 1 dataset and Experiment 2 dataset). We have employed a variation of Levene’s test, by extracting the residuals from the model, taking their absolute value, and squaring them (Glaser, 2006). The ANOVA between datasets for RT (F = 2.997, p = 0.085) and ACC (F = 1.011, p = 0.316) yielded no reliable results. Hence, the assumption of homoscedasticity was met for datasets’ residuals covariance. Finally, we tested the assumption of the normality of residuals, by generating QQ plots, which provide an estimation of where the standardised residuals lie with respect to normal quantiles (see Supplementary figure 4A-B).  Albeit with some slight deviation, the QQ plot for global RT suggested that the normality assumption was not violated, although some slight deviations were observed for ACC residuals. Nevertheless, violations of the normality assumption would not noticeably impact results if sample size exceeds 10 participants (Schmidt & Finan, 2018).

We also tested the main assumptions of linear mixed-effects models on the two hierarchical models we used to estimate the effects of congruency and the interaction between congruency and arousal on RT and ACC, with arousal, congruency and their interaction as fixed factors, and the random factors common to all models (participant nested into dataset). Again,  Shapiro-Wilk normality tests revealed the relationship between the variables and the RT (W = 0.967, p < 0.001) or the ACC (W = 0.909, p < 0.001) was not well described by a straight line. When checking whether the covariance of the residuals was equal across experiments, we observed again that the versioned Levene’s test ANOVA yielded no statistically reliable differences for dataset residuals in RT (F = 1.46, p = 0.23), but wasmidly reliable for ACC (F = 6.584, p = 0.011). Thus, the homoscedasticity assumption would be only partially fulfilled. Again, QQ plots suggested that probably RT residuals themselves are normally distributed, while some minor bias was observed in the ACC plot (see Supplementary figure 4C-D).

Finally, we tested the model used to assess RT conflict adaptation effects, in which arousal, congruency and previous congruency, and their three possible interactions were set as fixed factors. Testing the linearity of the data by plotting the R model’s residuals against the predictors and by implementing Shapiro-Wilk normality test yielded statistically reliable results (W =0.97, p < 0.001). Again, it simply means our dataset is not well described by a line, which would be consistent with the quadratic relationship found in RT in the results section. Testing the covariance of the residuals with the versioned Levene’s test showed reliability (F = 4.927, p = 0.027), while the QQ plot indicated that the residuals might not be normally distributed (see Supplementary figure 4E). In any case, given the sample size involved in our study, violation of the latter assumption should not have had a major impact on the results (Schmidt & Finan, 2018).

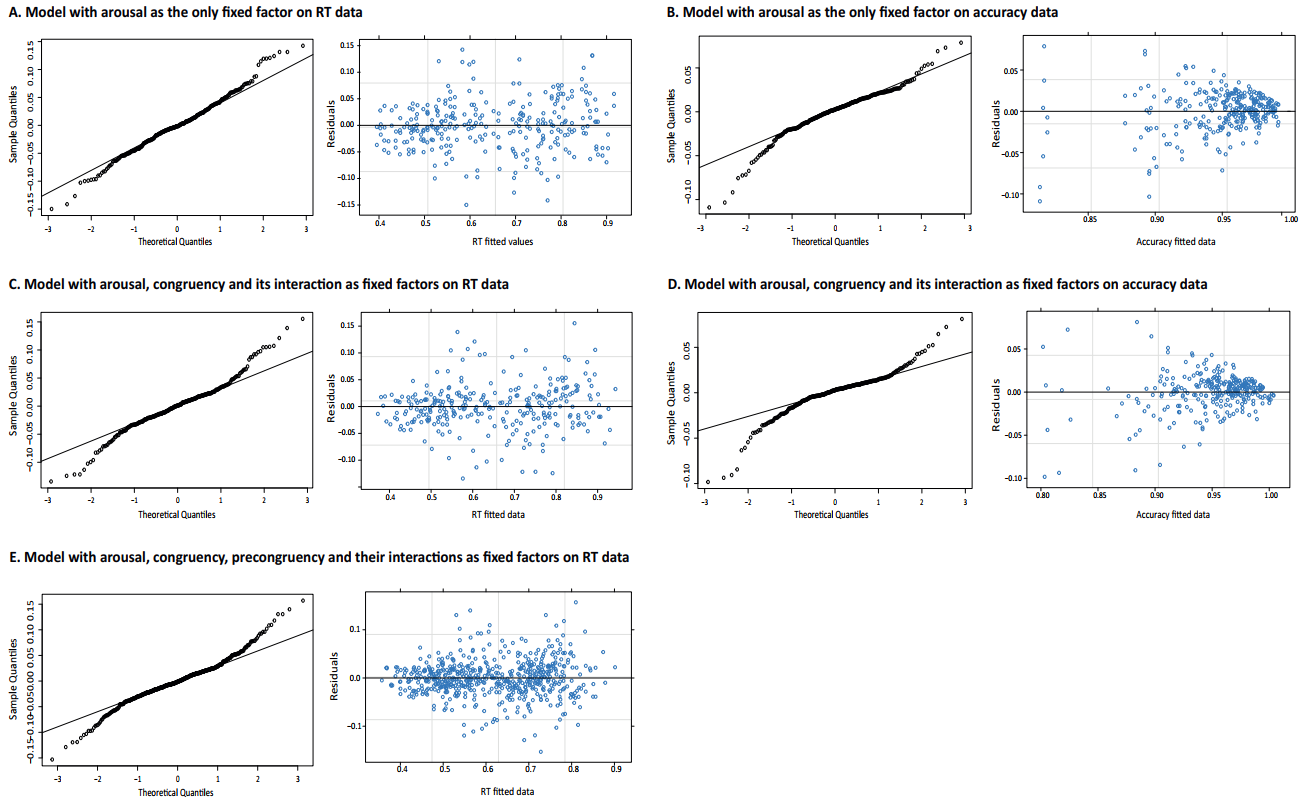
**Supplementary figure 4.** **Models for testing RT and accuracy hypotheses assumptions graphs. A-B)** Models with arousal as the only fixed factor, and participant nested into dataset as random factors (which are common to all models). For each model, the left panel shows the estimation of the normality of the residuals. Strong deviations from the provided line would indicate that the residuals are not normally distributed. The right panel represents fitted values versus model’s residuals. The absence of a concrete pattern suggests that the assumption has not been violated (Fox, 2008). **C-D)** Models with arousal, congruency and ‘arousal x congruency’ as fixed factor, to test hypotheses about conflict effect. **E)** Models with arousal, congruency, precongruency and their respective interactions as fixed factors, to test hypotheses about conflict adaptation effect.

***Models for testing differences in DDM parameters***

We tested the main assumptions of linear mixed-effects models on the five hierarchical models we used to estimate the effects of arousal on decision-making parameters, with arousal as the only fixed factor, and participant nested into dataset as random factors.

*Amplitude of automatic processes*

As with RT and accuracy, the relationship between arousal and the amplitude of automatic processes parameter could not be well described by a straight line (W = 0.9267, p < 0.001), which could be simply due to one variable being a sinusoidal function of the other, or a quadratic function, or even a straight line that changes slope at some point. However, testing homoscedasticity with the versioned Levene’s test validated the assumption (F = 2.961, p = 0.087), and the QQ plot for amplitude residuals did not show a strong deviation from the provided line (see Supplementary figure 5A).

*Time to peak of automatic processes*

The plot of the model’s residuals against the predictors, which does not show an absolutely random pattern (see Supplementary figure 5B), and the Shapiro-Wilk normality test on time to peak showed that the relationship between this parameter and arousal did not follow a straight line (W = 0.97421, p = 0.008). The variation of Levene’s test also yielded reliable results (F = 11.54, p < 0.001). Conversely, the QQ plot suggested that probably time to peak residuals are normally distributed  (see Supplementary figure 5B).

*Drift rate of controlled processes*

As with the previous parameters, the link between arousal and drift rate cannot be depicted by a straight line (W =0.97225, p = 0.005). According to Levene's test (F = 1.1787, p = 0.23) and the QQ plot, the assumptions of homoscedasticity and normality were both fulfilled (see Supplementary figure 5C).

*Boundary separation*

The assumption of linearity was validated by Shapiro-Wilk's test on boundary separation residuals (W = 0.989, p = 0.328) and the random distribution observed by plotting residuals against predictors (see Supplementary figure 5D), which is consistent with our results showing an increase in group mean boundary separation as arousal increases. Homoscedasticity (versioned Levene’s test = 2.992, p = 0.086) and linearity (QQ plot without deviations from the provided line) assumptions were also validated (see Supplementary figure 5D).

*Non-decision time*

In line with most variables here analysed, the relationship between arousal and non-decision time parameter could not be well described by a straight line (W = 0.956, p < 0.001). The variation of Levene’s test also yielded reliable results (F = 23.881, p < 0.001), while the QQ plot indicated that the residuals might not be normally distributed (see Supplementary figure 5E).

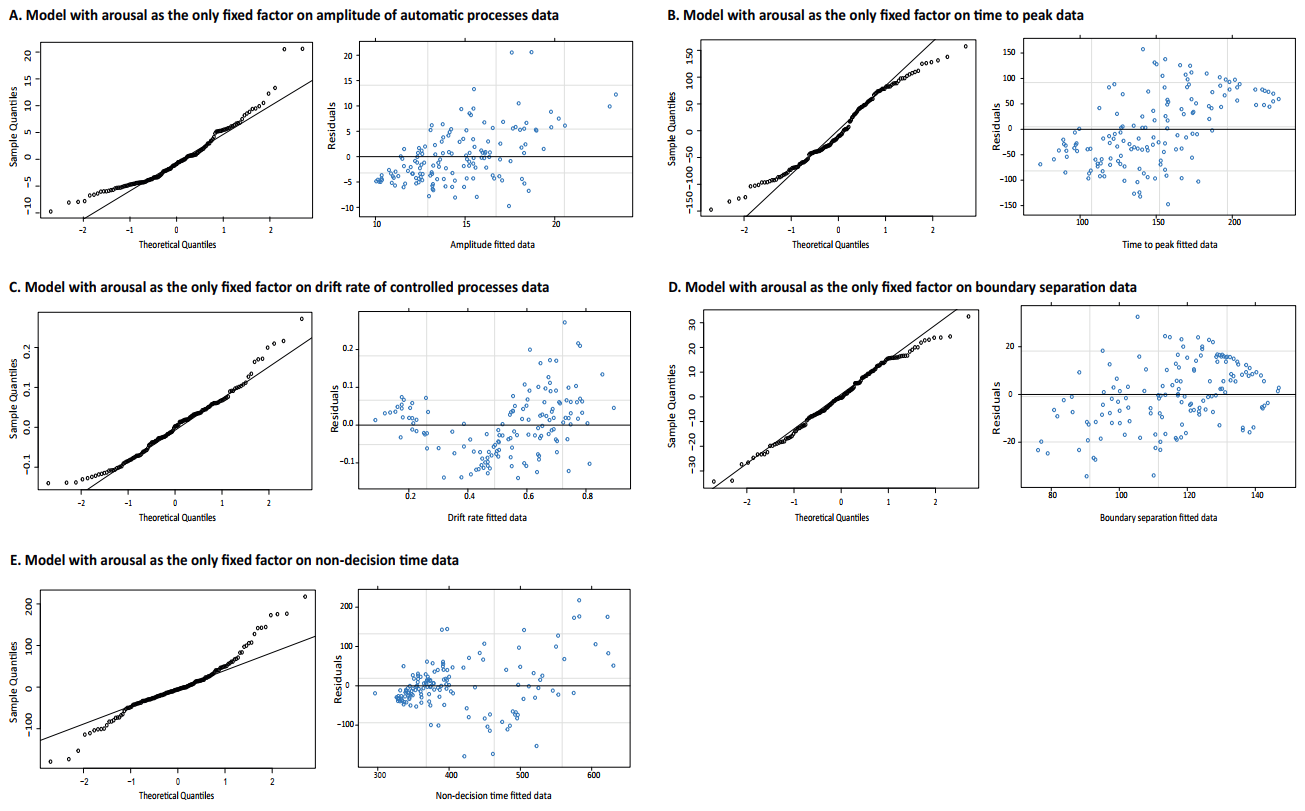
**Supplementary figure 5. Graphs on linear mixed-effects models assumptions for models testing the impact of arousal on DDM parameters: A) amplitude of automatic processes, B) time to peak of automatic processes, C) rate of evidence accumulation of controlled processes, D) separation between decision boundaries, and E) non-decision time.** For each model, the left panel shows the estimation of the normality of the residuals, while the right panel represents fitted values versus model’s residuals.

**Subject-by-subject mean heart rate and watts output during high arousal and high baseline conditions**

Individual heart rate (HR) and watts mean values per condition are provided in Supplementary table 1, to better illustrate the effective intensity of the induced physiological stress. After performing the incremental effort test, the watts output when each participant reached 80% of their maximum VO2 consumption (VO2 max) was used as target watts during the high arousal condition (referred to as “high arousal” in Supplementary table 1). Whereas, in the baseline condition (low-intensity exercise, reported in the Supplementary table 1 as "high baseline"), the target was 10% of the watts reached at VO2 max. Although 40 participants were initially recruited, data from one participant (subject 28 in Supplementary table 1) was discarded, as the HR achieved in the high intensity condition was considered too low (i.e., only 67% of his maximum HR, and over three standard deviations below the group average of 85% of HR max [SD = 5.06]).

**Supplementary table 1. Average HR and watts achieved by each participant in each experimental session, and average percentage relative to their maximum value.**

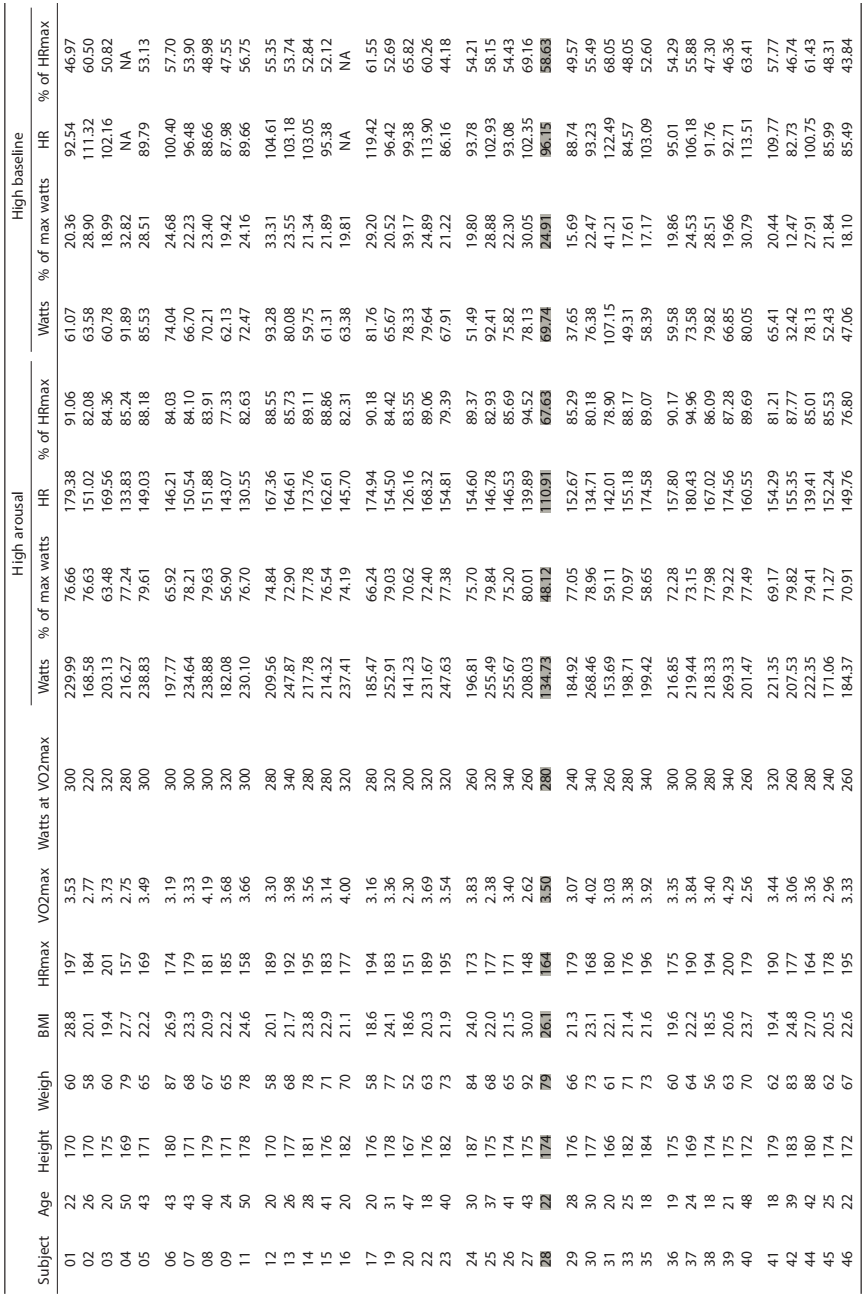
BMI: body mass index. HRmax: maximum heart rate. VO2max: maximum volume of oxygen consumption. The subject highlighted in grey was excluded from data analyses.
